# Thalamic contributions to psychosis susceptibility: Evidence from co-activation patterns accounting for intra-seed spatial variability (μCAPs)

**DOI:** 10.1101/2023.05.23.541896

**Authors:** Farnaz Delavari, Corrado Sandini, Nada Kojovic, Luigi F. Saccaro, Stephan Eliez, Dimitri Van De Ville, Thomas A. W. Bolton

**Author notes:** The symbol denotes shared authorship (Dimitri Van De Ville, Thomas A. W. Bolton).

## Abstract

The temporal variability of the thalamus in functional networks may provide valuable insights into the pathophysiology of schizophrenia. To address the complexity of the role of the thalamic nuclei in psychosis, we introduced micro-co-activation patterns (μCAPs) by employing this method on the human genetic model of schizophrenia 22q11.2 deletion syndrome (22q11.2DS). Participants underwent resting-state functional MRI and a data-driven iterative process resulting in the identification of six whole-brain μCAPs with specific activity patterns within the thalamus. Unlike conventional methods, μCAPs extract dynamic spatial patterns that reveal partially overlapping and non-mutually exclusive functional subparts. Thus, the μCAPs method detects finer foci of activity within the initial seed region, retaining valuable and clinically relevant temporal and spatial information. We found that a μCAP showing co-activation of the mediodorsal thalamus with brain-wide cortical regions was significantly less frequent in patients with 22q11.2DS, and its occurrence negatively correlated with the severity of positive psychotic symptoms. Additionally, the activity within the auditory-visual cortex and their respective geniculate nuclei were expressed in two different μCAPs. One of these auditory-visual μCAPs co-activated with salience areas, while the other co-activated with the default mode network (DMN). A significant shift of occurrence from the salience+visuo-auditory-thalamus to the DMN+visuo-auditory-thalamus μCAP was observed in patients with 22q11.2DS. Thus, our findings support existing research on the gatekeeping role of the thalamus for sensory information in the pathophysiology of psychosis and revisit the evidence of geniculate nuclei hyperconnectivity with the audio-visual cortex in 22q11.2DS in the context of dynamic functional connectivity as specific hyper-occurrence of these circuits with the task negative brain networks.

## Introduction

Resting-state functional magnetic resonance imaging (RS-fMRI) indirectly quantifies neuronal activity by measuring the blood oxygenation level-dependent (BOLD) signal [1]. Studying brain function using RS-fMRI has become pivotal in shaping our understanding of the neural mechanisms that underlie neurological and psychiatric disorders [2, 3]. Lately, temporal fluctuations in brain activity and reconfigurations in cross-regional functional interactions have been gaining attention, and have been studied with various quantitative and modeling approaches [4-7], collectively summarized under the dynamic functional connectivity (dFC) umbrella term (see [8, 9] for reviews). Taking into account time-resolved features of resting state brain function has further advanced our understanding of several brain disorders [10, 11].

Seed-based functional connectivity (FC) is a longstanding approach in which statistical interdependence (Pearson correlation) of activity of a seed region-of-interest (ROI) with the rest of the brain is quantified [12]. It only reflects average FC over the course of an fMRI recording. Co-activation pattern (CAP) analysis is a time-resolved extension. During CAP analysis, fMRI volumes for which the seed signal exceeds a threshold are selected. While simple averaging of these volumes would be a proxy for seed-based FC, clustering of the selected fMRI volumes yields CAPs, i.e., states of seed co-(de)activation with the rest of the brain that recur over time and across subjects and can then be analyzed in terms of their spatiotemporal dynamics [13-15].

CAP analysis is a versatile tool that has been successfully applied, not only to RS-fMRI in order to study pathological brain conditions [16-19], but also in task-based or neurofeedback settings [20, 21]. However, recent work has evidenced the remaining limitations of the approach: for example, similar CAP-based metrics could be derived from a stationary null model, evidencing that the interpretation of CAP findings in terms of temporal dynamics should remain cautious [22]. Another current limitation of CAP analysis lies in the implicit assumption that the seed region used for the selection of fMRI volumes to cluster is functionally homogeneous. In the present work, we propose an extension of the CAP methodology that accounts for intra-seed spatial variability. As a key example, we focus on the thalamus as a seed region to illustrate shortcomings of conventional CAP analysis and how these are resolved with the proposed framework.

The thalamus is involved in a wide range of functions including motor control, and attention, salience, and sensory processing [23]. It acts as a relay station for information coming in from the senses and going out to the rest of the brain [24]. The thalamus is also involved in regulating states of consciousness, sleep, and wakefulness [25]. It is thought to play a key role in the development of several neurological and psychiatric disorders, including schizophrenia and psychosis spectrum disorders [26, 27]. Indeed, aberrant processing of the saliency of sensory stimuli has been proposed as a pathophysiological model of psychosis, postulating that psychotic behavior is driven by the inappropriate processing of sensory information that would normally be deemed irrelevant, due to disproportionate assignment of salience and importance to this information, which may then lead to pathological orientation of attention and, ultimately, to delusional symptoms and behavior[28]. FMRI studies of the thalamus allow for non-invasive investigation of the neural networks that underlie these functions. For example, in agreement with the role of the thalamus in salience and sensory processing, RS-fMRI studies have revealed important information about how disturbed thalamic functional connectivity gives rise to psychotic symptoms [29, 30]. DFC RS-fMRI investigations have further demonstrated the clinical relevance of thalamic FC. For example, patients with schizophrenia show dynamic hyperconnectivity of the thalamus with auditory and visual networks [31, 32]. Overall, temporally variable involvement of the thalamus within different functional networks thus contributes to aberrant symptomatology, which positions this brain structure as an ideal candidate for CAP analysis.

However, the thalamus is also a complex, multi-faceted area divided into several subnuclei, including the dorsal thalamus, ventral thalamus, and the intralaminar nuclei [33, 34]. Studying the subnuclei of the thalamus is essential for understanding the neural mechanisms underlying several brain functions and how they may relate. The dorsal thalamus is involved in the processing of somatosensory information [35], while the ventral posterior thalamus plays a critical role in relaying visual and auditory information to the cerebral cortex [36]. The intralaminar nuclei are involved in the regulation of arousal and attention [37]. Dysfunction of certain circuits involving these nuclei have been linked to a range of symptoms in schizophrenia [38]. However, the anatomical segmentation of the thalamus is not necessarily reflective of its functional organization. As the thalamus is often characterized as a hub for diverse connections within the human brain, it has been a candidate region for methods that employ functional-based segmentations. Yet, these methods often disregard the dynamic nature of thalamic connectivity, despite its aforementioned clinical relevance [26]. Furthermore, the evidence on multiple functionality of hubs within the thalamus [39] suggests that there is no one-to-one correspondence between the subnuclei of the thalamus and the cortex and subcortex. To retain this type of physiological information, an accurate analysis of thalamic activity dynamics must allow for partially overlapping and non-mutually exclusive functional subparts.

Here, we introduce an extension of CAP analysis to extract so-called micro-CAPs (μCAPs). As opposed to conventional CAPs, μCAPs allow for a spatial pattern within the seed, thus potentially revealing functional subdivisions with regards to its dynamic functional interactions with the rest of the brain. By using this data-driven approach, we aim to avoid the limitations of anatomical prior knowledge and to study the functional organization of the thalamus in a more comprehensive manner. We assess the physiological pertinence of μCAPs in terms of thalamic nuclei connections to the whole brain. Furthermore, we illustrate how, compared to conventional CAPs, μCAPs provide new insights into the function of the thalamus and its contribution to psychopathology. To do so, we study a population of adults with 22q11.2 deletion syndrome (22q11.2DS). Affecting up to 1 in 3000 live births, 22q11.2DS is the most common human microdeletion [40]. Even though it leads to a fortyfold increase in the risk of developing psychosis, early or preventive interventions for psychosis are still not explicit in this population. This could be in part attributed to the lack of objective biomarkers [41, 42]. In pursuit of identifying neuroimaging biomarkers of susceptibility to psychosis in patients with 22q11.2DS and other at-risk populations, we propose that μCAPs can more robustly explain the diagnostic differences between healthy controls and patients with 22q11.2DS, as well as the severity of positive psychotic symptoms in this latter population.

## Materials and Methods

### Participants

Overall, 106 adult participants (age span 18-35) underwent structural and functional brain imaging and were included in the analysis. Participants were part of an ongoing cohort of 22q11.2DS patients and matched healthy controls (HCs) recruited in Geneva, Switzerland. The sample included 50 individuals with confirmed diagnosis of 22q11.2DS, and 56 HCs. The latter were screened for present or past medical history of neurological or psychiatric disorders and were recruited from siblings of deletion carriers or through the Geneva state school system. The research protocols were approved by the Institutional Review Board of Geneva University School of Medicine. Written informed consent was obtained from all participants.

A comprehensive clinical assessment with an expert psychiatrist (SE), including a semi-structured clinical interview [43], and the Positive and Negative Syndrome Scale (PANSS) [44], was conducted on participants with 22q11.2DS. The positive subscale of the PANSS is measuring the severity of positive psychotic symptoms (Delusions, Conceptual disorganization, Hallucinatory behavior, Excitement, Grandiosity, Suspiciousness, Hostility), and is the focus of the analysis in what follows in this study. **Table 1** summarizes the demographic and clinical characteristics of the participants included in the study.

**Table 1.** Demographic and clinical information about the analyzed subject populations. When applicable, mean and standard deviation are reported. ^a^: Paired t-test significance assessment, ^b^: χ2 test significance assessment, ^***^: *p* < 0.001.

| Variable Name | 22q11.2DS | HC | P-value |
| --- | --- | --- | --- |
| Number | 50 | 56 | NA |
| Age (average (SD)) | 22.8 (3.1) | 22.7 (3.4) | 0.87 <sup>a</sup> |
| Sex (female percentage) | 56.0% | 50.0% | 0.53 <sup>b</sup> |
| Schizophrenia (positive percentage) | 3.5% | NA | NA |
| IQ (average (SD)) | 108.6 (18.7) | 122.9 (12.9) | <0.001 *** <sup>a</sup> |
| PANSS (average (SD)) | 38.6 (10.1) | NA | NA |

### Magnetic resonance imaging (MRI) and preprocessing

MRI brain images were acquired on a Siemens Trio (Siemens Healthineers) 3T scanner. Structural images consisted of a T1-weighted sequence with 0.86 × 0.86 × 1.1 mm^3^ volumetric resolution (192 coronal slices, repetition time = 2500 ms, echo time = 3 ms, acquisition matrix = 224 × 256, field of view = 22 cm^2^, flip angle = 8°). RS-fMRI scans were obtained by asking participants to fixate a white cross on a black screen and stay awake, consisted of a T2-weighted sequence (200 frames, voxel size = 1.84 × 1.84 × 3.2 mm^3^, repetition time = 2400 ms) and lasted for 8 minutes (equivalent to a total of 200 functional volumes).

The acquired structural and functional images were preprocessed with a previously described and published in-house pipeline [45]. Briefly, it uses SPM12 (Wellcome Trust Centre for Neuroimaging, London, UK; http://www.fil.ion.ucl.ac.uk/spm), Data Processing Assistant for Resting-State fMRI (DPARSF [46]), and Individual Brain Atlases using Statistical Parametric Mapping (IBASPM [47]). We performed realignment of functional images, spatial smoothing with an isotropic Gaussian kernel of 3 mm full width at half maximum (FWHM), and co-registration of structural scans to the functional mean. Segmentation of the structural images was done using SPM12’s segmentation algorithm [48] and Diffeomorphic Anatomical Registration using Exponential Lie algebra (DARTEL [49]) was used to compute a study-specific template. The first 5 functional scans were excluded from the analysis to account for magnetization equilibration. The time series were linearly detrended and translational and rotational movement parameters as well as global, mean white matter and mean cerebrospinal fluid signals were regressed out from the BOLD time series using the warped DPARSF [46] tissue masks. The time series were subsequently filtered with a bandwidth of 0.01 Hz to 0.1 Hz. After preprocessing, all functional images were warped to the DARTEL space, and then to MNI space to create population-based results. Extended motion correction through scrubbing of frames with high framewise displacement (FD > 0.3 mm [50]) is included within the μCAPs algorithm to exclude them from the analysis.

### Extraction of μCAPs

We outline the key steps regarding the extraction of μCAPs. The code for this process is available at: https://github.com/MIPLabCH/mCAP. The whole process is illustrated in **Figure 1**. Outputs from this new algorithm were compared to conventional CAPs extracted with the *TbCAPs* toolbox [51], as further detailed in later subsections.

**Figure 1.**
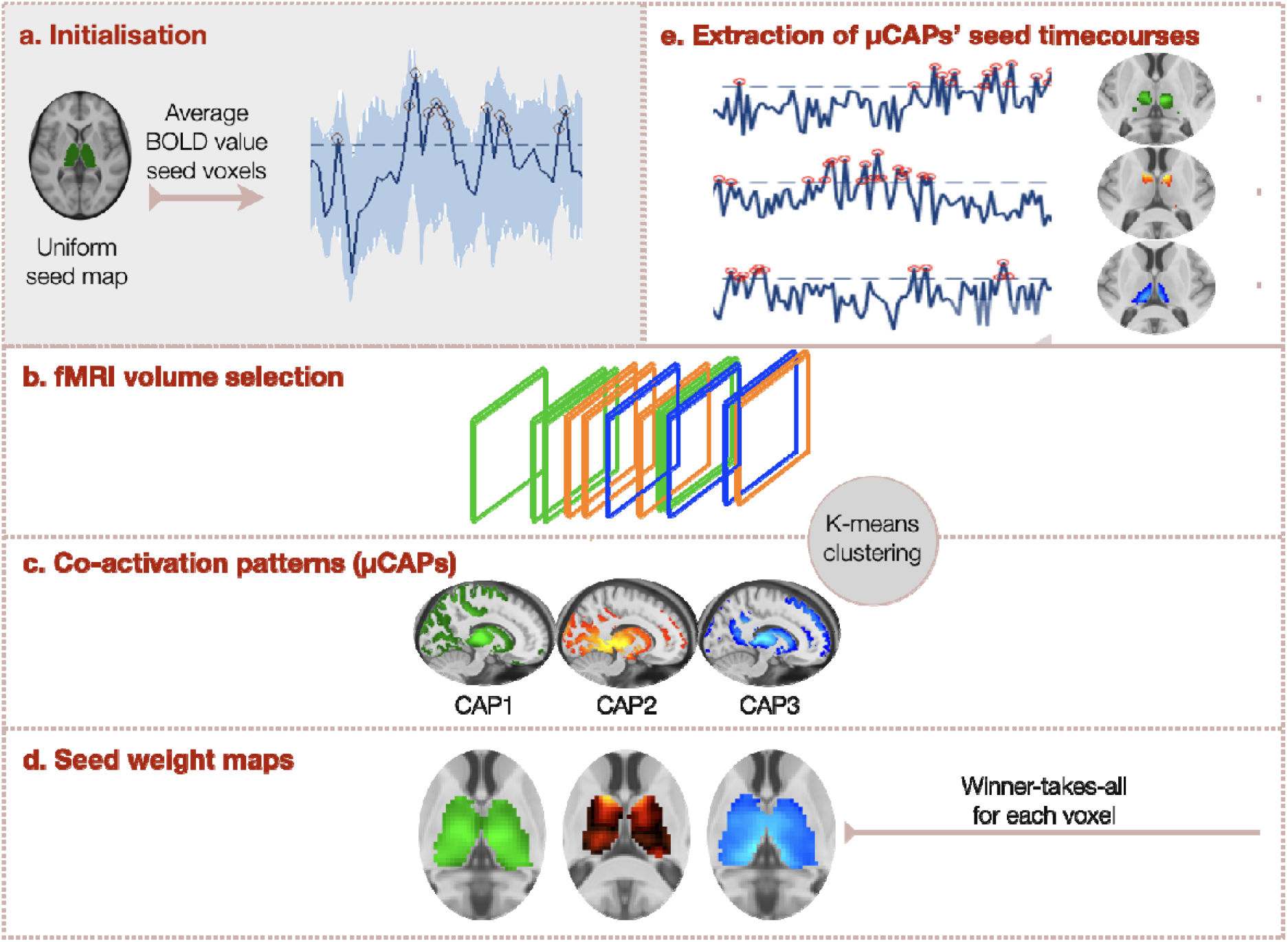
Flowchart of the μCAPs processing pipeline. (**a**-**b**) Initially, the seed region is considered uniform (top left, shown in green), and a Z-scored time course of seed activity is created and thresholded at 0.5 to extract moments of significant seed activity. (**c**) The selected fMRI volumes undergo K-means clustering into μCAPs, and (**d**) The associated weight maps within the seed region are extracted. The weight maps are processed to retain only the maximal seed signal for each voxel, and (**e**) new seed time courses are created and thresholded at 1 to retain a new set of fMRI volumes. The process is repeated until convergence (see **Supplementary Materials** – Figure S3).

#### a) Notations and initialization

Let us consider the BOLD fMRI data for all concatenated subjects as a matrix *X* of size *V* × *T* containing the time-dependent fMRI volumes as its columns, where *V* is the number of voxels entering the analysis and *T* the total number of available time points across all subjects. We choose a seed ROI as the subset of voxels *S*. The aim of the method will be to identify *K* μCAPs where each μCAP is also associated with a weight map for the seed ROI: *S*_*k*_(*v*), *v* ∈ *S, k =* 1, …, *K*. To initialize the algorithm, we put *S*_*k*_(*v*) = 1, for all *k* and *v*.

#### b) Selection of frames

We compute, for each seed weight map *S*_*k*_, the corresponding time course:

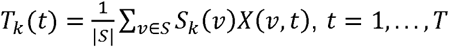, where *t* is the currently considered time point. The time course is then further Z-scored over time, as in conventional CAP analysis.

Given the initialization above, all *K* time courses will be identical for the first iteration. In later stages of the algorithm, they then highlight the temporal activity of different seed patterns.

For each seed map, fMRI volumes are tagged as “selected” if the Z-scored time course exceeds a certain threshold. Unless otherwise stated, for the first iteration, we choose this threshold to be 0.5 to maximize sensitivity. For subsequent iterations, the threshold is set to 1.0, a value within the range of typical choices for conventional CAP analyses [21, 51].

#### c) Clustering

Selected fMRI volumes for all seed maps are then be clustered using the K-means method for *K* clusters, based on a 1-correlation distance metric. The process of finding the optimal *K* is described below. For each cluster, the representative centroid is obtained as the average over the frames assigned to it. These centroids are called micro co-activation patterns (μCAPs, **Figure 1b**), denoted *C*_*k*_(*v)* with *v* = 1, …, *V*.

#### d) Update of seed weight maps

To determine the new seed weight maps, we first restrict each of the μCAPs to the seed region (**Figure 1c**): *S*_*k*_(*v*) = *C*_*k*_(*v*), for *v* ∈ *S*. We then keep, for each voxel, only the largest μCAP value; *i*.*e*., a “winner-takes-all” approach. Values for the other μCAPs are set to zero (**Figure 1d**). The algorithm then iterates and goes to step b) again.

Convergence is monitored by keeping track of the distance between the μCAP seed maps across iterations. The distance between seed patterns within the μCAPs of two successive iterations is computed by first matching them using the Hungarian algorithm [52], and then computing the average cosine distance across pairs. The threshold for convergence was set at 0.005. The first iteration with the initial uniform seed pattern was left out of the convergence assessment.

### Thalamic μCAPs and application to 22q11.2DS

On the 22q11.2DS cohort experimental data, we considered the bilateral thalamus from the Automated Anatomical Labeling (AAL) atlas [53] as the seed region. CAPs were derived with *TbCAPs* with a frame selection threshold of 1, and μCAPs were extracted using the proposed algorithm. In both cases, frames associated to excessive framewise displacement (FD > 0.3 mm) were discarded.

The optimal *K* for experimental data was chosen using two different criteria based on a test – training K-means clustering process. In this process, we performed a 20-fold random 50-50 split of subjects into training and testing subsamples. For the first criterion, the training frames went through K-means clustering to produce the *K* different centroids. The distance of the test frames to the *K* training centroids (assigning each test frame to its closest training centroid) was calculated. The cluster with the highest sum of distances of test frames was chosen as the worst-fit cluster and it was monitored across a range of candidate *K* values (2 to 10). As expected, the sum of distances decreases as *K* increases. However, a local drop in the sum of distances for the worst-fit cluster indicated an optimal fit for a candidate *K*. For the second criterion, both test and training frames went through K-means clustering to produce *K* different centroids each. Then, the *K* clusters produced by test and training frames were matched using the Hungarian algorithm and the average of *K* distances across the matched centroids was calculated for a range of candidate *K* values (4 to 8). The lowest distance of matched centroids would indicate maximal similarity of results across test and training frames and therefore, stable K-means results. In our dataset, both criteria converged on the same optimal solution. This process is described in details in the **Supplementary Materials**.

Obtained CAPs and μCAPs were analyzed in terms of their spatial patterns of seed co-(de)activation, and qualitatively contrasted across methods. To quantitatively assess clinical relevance, for each participant, we considered the number of frames assigned to each (μ)CAP as a metric of interest. These imaging metrics were correlated with the T-score of the PANSS positive symptom subscale. Results were corrected for multiple comparisons using the false discovery rate (Benjamini Hochberg) method [54].

## Results

To validate and illustrate the benefits of μCAPs over traditional CAPs, we generated simulated data with characteristic spatiotemporal properties the results are depicted in **Supplementary Materials** (**Figure S1**).

### Characterization of μCAPs on experimental data

On experimental data, 86.46% of the frames had an FD lower than 0.3 mm and thus contributed to the analysis. However, the average FD after scrubbing significantly differed across groups (mean (SD) for HC=0.11(0.03), mean (SD) for 22q11.2DS=0.14(0.03), p<0.001). Additionally, as expected, the average IQ was significantly different across groups. Therefore, as a supplementary analysis, average FD after scrubbing and IQ were regressed out as covariates of no interest when assessing group differences (see **Supplementary Materials** for further details). The groups were matched on sex and age. The demographic information is displayed in **Table 1**.

Following the test – training K-means clustering process, both criteria converged on *K* = 6 as an optimal solution for number of clusters for our dataset. Detailed information is displayed in the **Supplementary Materials** (**Figure S2**).

μCAPs are introduced in decreasing order of temporal occurrences. Sagittal, coronal, and axial slices are displayed for each μCAP in **Figure 3**, while **Table 2** summarizes brain co-(de)activations as well as the within-thalamus patterns captured by each μCAP. **Figure 4** (left half) more comprehensively displays the seed patterns obtained for each μCAP.

**Figure 3.**
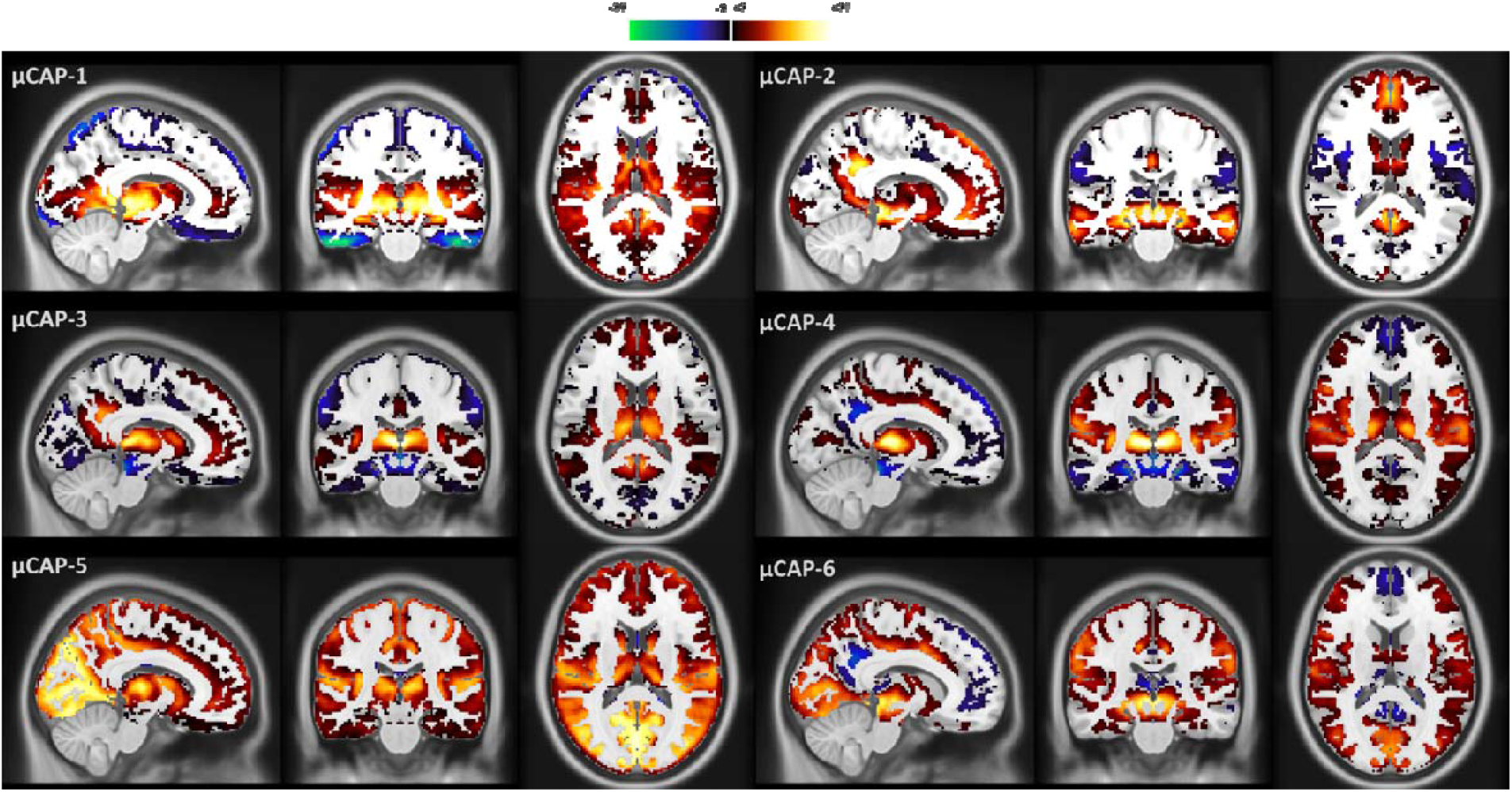
The six μCAPs extracted from experimental data. Spatial T-score maps calculated, for each voxel, as the average value for frames assigned to each activation pattern (μCAP) divided by the standard deviation of the voxels across those frames, multiplied by the square root of the number of frames. Thresholded at 3 corresponding to *p* < 0.01. μCAPS are displayed in descending order of temporal occurrences (respectively [mean±SD]: 10.9±6.6, 10.4±4.7, 10.2±5.1, 10.1±5.3, 9.8±7.1, 8.7±4.6 frames across subjects).

**Table 2.** Summary of μCAPs results. For all six μCAPs, foci of co-(de)activation with the thalamus, as well as exact patterns found within the initial thalamic seed region, are reported. VTA: ventral tegmental area; SN: substantia nigra; SC: superior colliculus; dlPFC: dorsolateral prefrontal cortex; dACC: dorsal anterior cingulate cortex; DMN: default mode network; mPFC: medial prefrontal cortex; PCC: posterior cingulate cortex; IC: inferior colliculus.

| μCAP | Within-thalamus pattern | Co-(de)activation with thalamus |
| --- | --- | --- |
| 1 | Anteroventral – Ventral posterolateral – Pulvinar – Lateral and medial geniculate nuclei – Ventral midline | + Primary visual cortex, middle and superior temporal cortex, insula, amygdala, hippocampus, putamen, midbrain structures (VTA, SN, SC)<br>- dlPFC, dACC |
| 2 | Ventral midline thalamus | + DMN (mPFC, PCC, precuneus, and angular gyri), middle temporal cortex, hippocampus, amygdala, midbrain structures (VTA, SN, SC, and IC)<br>- Insula and juxta positional lobule cortex |
| 3 | Dorsal midline thalamus | + DMN (mPFC, PCC, precuneus, and angular gyri), posterior insula, putamen, caudate, nucleus accumbens<br>- Hippocampus, amygdala, and midbrain structures |
| 4 | Mediodorsal | + Insula, juxta positional lobule cortex, dorsal anterior cingulate cortex (dACC), dlPFC, posterior parietal cortex, pre-central and post-central cortex, putamen<br>- DMN, hippocampus, amygdala, and midbrain structures. |
| 5 | Mediodorsal – Inferior pulvinar – Lateral and medial geniculate nuclei | + Occipital lobe, superior temporal cortex, insula, pre-central and post-central cortices, dACC, mPFC, superior frontal cortex, dlPFC, caudate, nucleus accumbens |
| 6 | Anteroventral – Ventral midline thalamus – Medial pulvinar – Lateral and medial geniculate nuclei | + Occipital cortex, superior temporal cortex, dACC, dlPFC, juxta positional lobule cortex, hippocampus, amygdala, midbrain structures (VTA, SN, SC, and IC)<br>- DMN (mPFC, PCC, precuneus, and angular gyri) |

**Figure 4:**
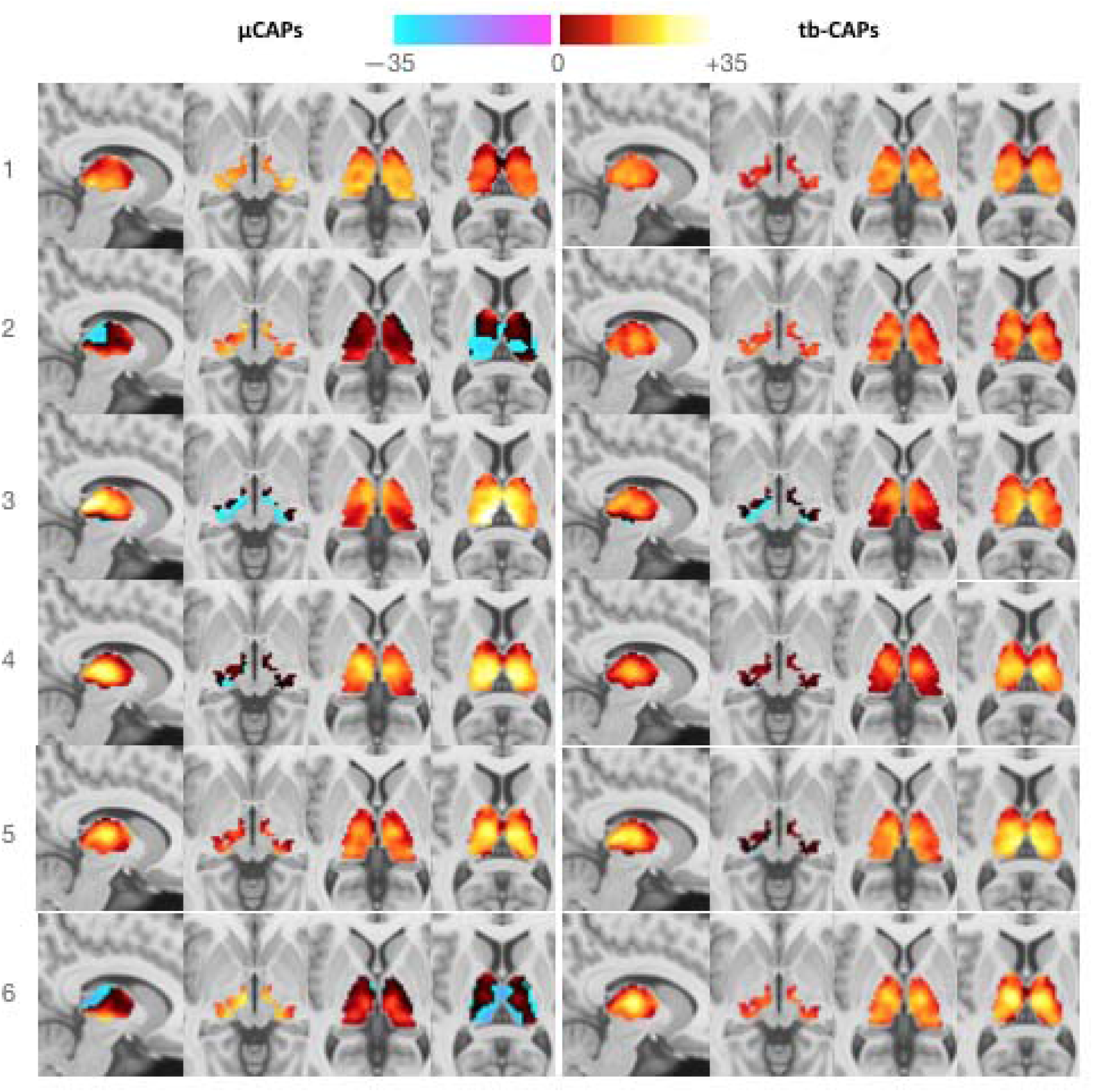
Seed patterns obtained when extracting μCAPs and conventional CAPs. μCAPs 2 and 6 include negative seed signals. Conventional CAP seed patterns are more spatially homogeneous.

μCAP1 captured the co-activation of several thalamic subnuclei with visual and auditory cortices, along with dopaminergic and limbic subcortical areas. μCAP2 and μCAP3 both showed co-activation of the midline thalamus with the default mode network (DMN). However, μCAP2 also displayed co-activation with limbic and midbrain regions, whereas μCAP3 instead included co-deactivations *(i*.*e*., negative-valued signal) of these areas, as well as ventral striatum co-activation. Moreover, the dorsal midline thalamus was active in μCAP3 whereas it was deactive in μCAP2. Of note, μCAP2 and μCAP6 were the only ones featuring both negative and positive values within the initial seed region. μCAP4 captured a pattern of salience and executive network co-activation along with deactivation of the DMN. Within the thalamus, the mediodorsal thalamus was consistently active for μCAP4. μCAP5 captured a diverse cortical pattern in which prefrontal, visual, auditory, and somatosensory cortices, as well as the ventral striatum, co-activated with selected thalamic subnuclei. The parahippocampus, hippocampus, and amygdala were absent from this pattern. Within μCAP6, the dorsolateral prefrontal cortex (dlPFC) and dorsal anterior cingulate cortex (dACC) were co-active with the thalamus, whereas the medial PFC as well as other parts of the DMN co-deactivated. Moreover, co-activations were also seen in the visual, auditory, and juxta positional cortices, as well as in limbic and dopaminergic regions.

### Comparison between μCAPs and conventional CAPs

To investigate differences in the information represented by μCAPs as opposed to conventional CAPs, we also extracted the latter on the same dataset. The six conventional CAPs are depicted in **Figure S4** (see **Supplementary Materials**). As expected, the patterns of negative value within the seed observed in some μCAPs were not present in this case (see also **Figure 4**, right half).

**Figure 5a** shows pairwise correlation values for the six CAPs derived with both methods, matched according to the Hungarian algorithm [44] based on cosine distance. High correlations, and therefore good matches, were detected for μCAPs 1, 3, and 4. For μCAPs 2, 5 and 6, lower correlations were observed.

**Figure 5.**
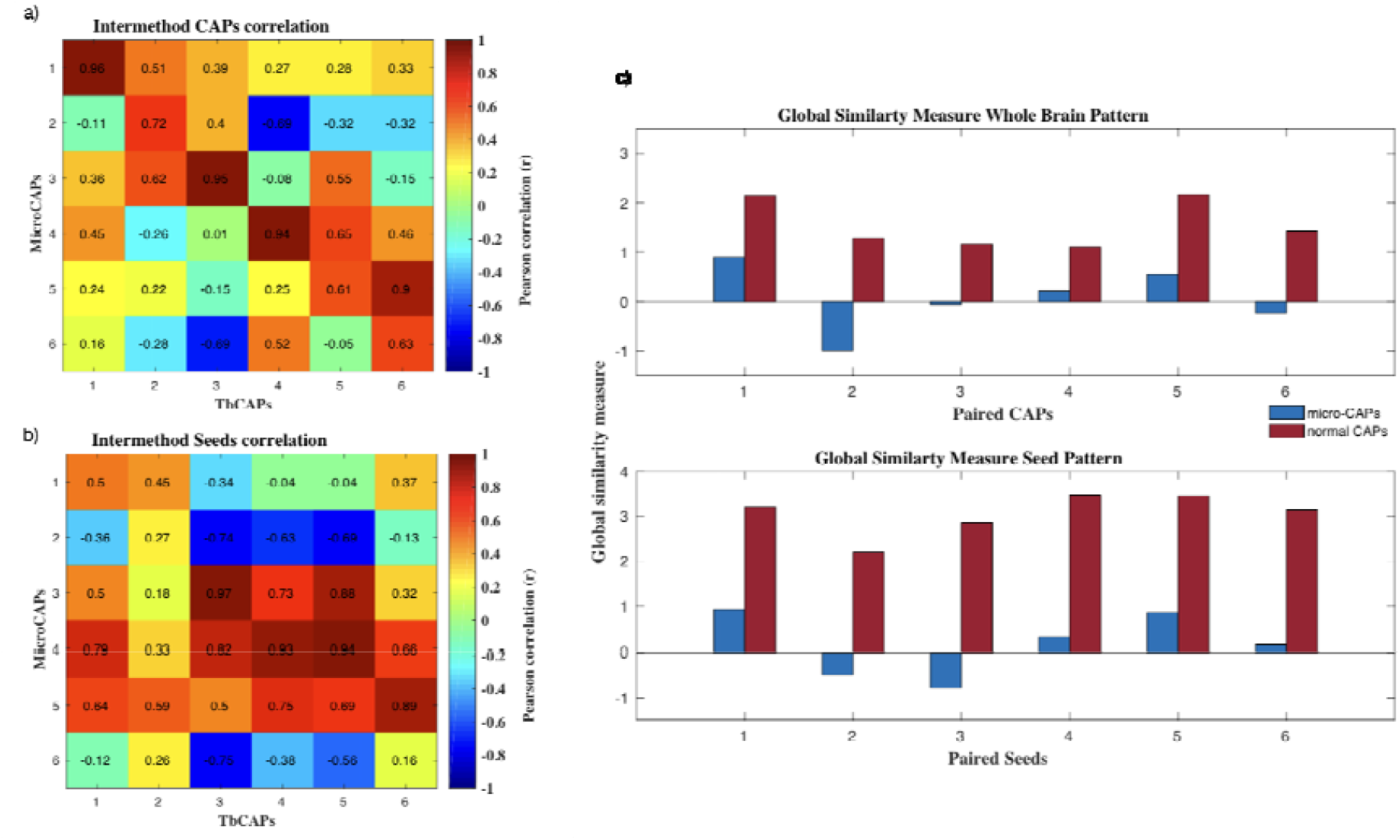
Co-activation pattern similarity across methods. Pairwise spatial correlations between CAPs obtained using the conventional approach and the proposed algorithm are displayed for whole-brain patterns (**a**) and within-seed patterns (**b**). The sums of correlation values to a given (μ)CAP are also reported (**c**).

Applying the same matching logic specifically to the seed patterns **(Figure 4b)**, similarity to the thalamic activation patterns found in conventional CAPs was consistently low for μCAPs 2 and 6. To assess spatial diversity of the retrieved patterns, we additionally computed Pearson’s correlation coefficient for each CAP with the other five CAPs obtained from the same method. **Figure 4c** shows the sum of the 5 correlation values for each CAP obtained from μCAPs in red and from conventional CAPs in green. This global similarity measure was consistently lower for the whole-brain patterns as well as the within-thalamus patterns obtained from μCAPs.

**Table 3** shows the number of frames that were assigned to a certain CAP only using μCAPs, only using conventional CAPs, and the number of frames that were assigned to a certain CAP in both methods.

**Table 3.** Frame selection across methods.

| CAP number<br>(Matched with Hungarian algorithm across methods) | Number of frames |  |  | Correlation of the CAPs across methods |
| --- | --- | --- | --- | --- |
| | Captured by μCAPs only | Captured by TbCAPs only | Captured by both methods | |
| 1 | 762 | 63 | 394 | 0.96 |
| 2 | 920 | 229 | 183 | 0.72 |
| 3 | 774 | 50 | 304 | 0.95 |
| 4 | 800 | 61 | 270 | 0.94 |
| 5 | 925 | 347 | 115 | 0.61 |
| 6 | 820 | 371 | 102 | 0.63 |

μCAPs consistently captured more frames when compared to conventional CAPs. As expected, for the matched CAPs for which correlation was low, fewer mutual frames were selected across methods. We contrasted temporal occurrence for each (μ)CAP between patients with 22q11.2DS and HCs (**Table 4**).

**Table 4.**
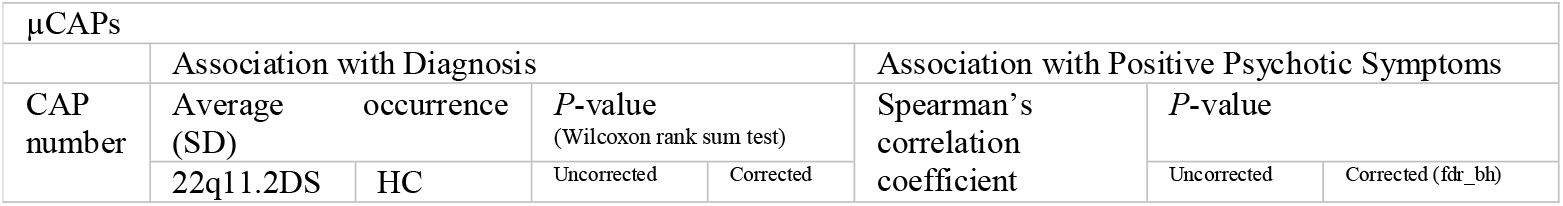
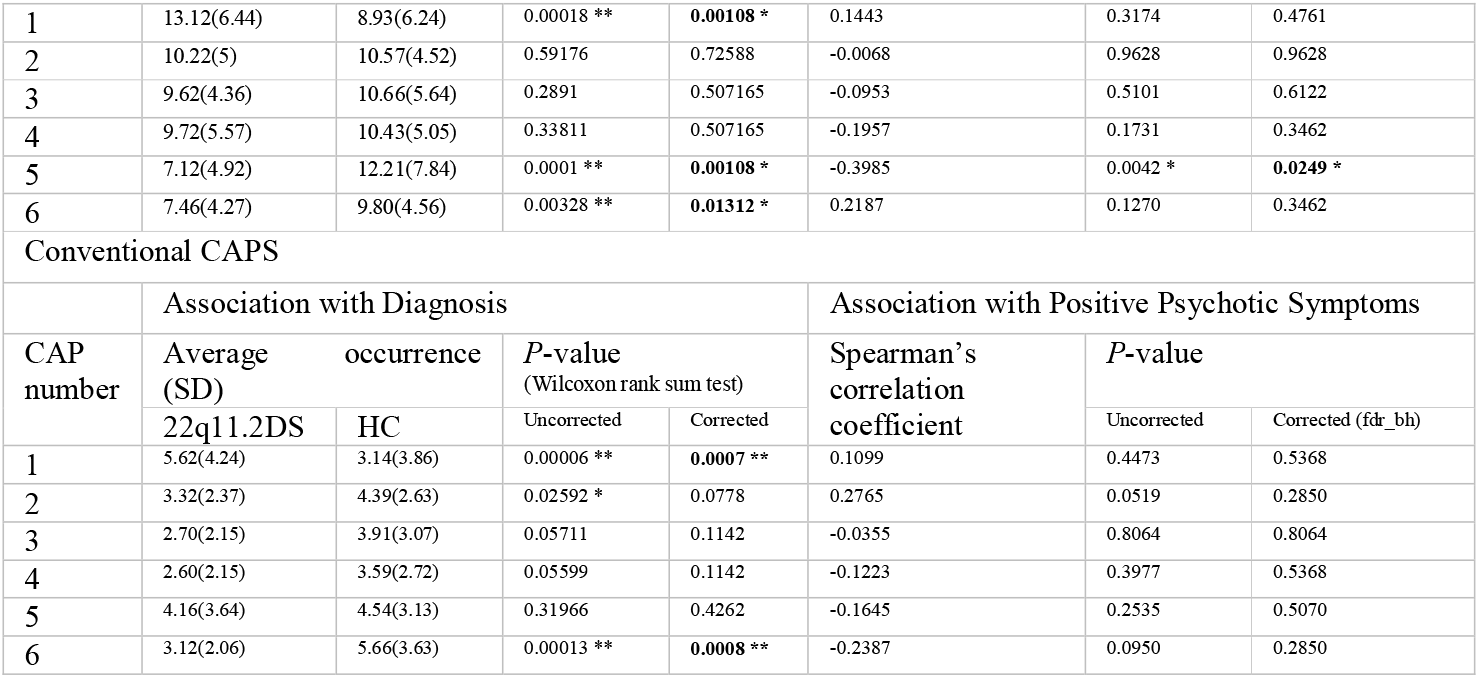
Association of temporal occurrences with diagnosis and psychopathology. Significance outcomes upon correction are highlighted in bold (^*^: *p*<0.05, ^**^: *p*<0.005).

Among μCAPs, μCAP1 occurred significantly more frequently in patients with 22q11.2DS. μCAPs 5 and 6 occurred significantly less in patients with 22q11.2DS. Among the standard CAPs, CAPs 1 and 6 showed higher and lower occurrences in participants with 22q11.2DS, respectively. To assess whether the larger amount of fMRI volumes assigned to μCAPs holds meaningful clinical information, temporal dynamics of the (μ)CAPs were correlated with positive psychotic symptoms of the 22q11.2DS patients. μCAP5 showed significantly lower occurrences in patients with higher scores for positive psychotic symptoms.

The μCAP1 and μCAP6 had a similar within-the-seed pattern (r=0.68), but their differences in occurrences between healthy controls and patients with 22q11.2DS were opposite. Therefore, we performed a post-hoc analysis considering the ratio of the occurrence μCAP6 over μCAP1, to assess whether the within-subject balance-shift of expression to μCAP6 is a significant characteristic. The median of this ratio was 1.17 in HC and 0.56 in patients with 22q11.2DS (Wilcoxon rank test, P-value<0.001). **Figure S5** shows the receiver operating characteristic (ROC) curve when using this single measure for detecting patients with 22q11.2DS diagnosis, which resulted in an area under the curve (AUC) of 0.75.

## Discussion

Here, we introduced an extension of CAP analysis by accounting for within-seed spatial variability. We demonstrated that the proposed iterative algorithm leads to a different within-seed weight map for each μCAP that reflects the heterogeneous engagement of the seed with a certain whole-brain pattern. Moreover, we showed that in comparison to conventional CAPs, frame selection is improved and, therefore, whole-brain patterns become not only more detailed, but also more relevant for applications, such as illustrated on a clinical population with 22q11.2DS. In short, we argue that the improved temporal representation of each μCAP allows for better behavioral and clinical phenotyping.

In the following sections, we first discuss the relationships between the exposed whole-brain pattern of co-(de)activation and within-thalamus activity in light of the existing literature. Likewise, we examine the implications of altered μCAP temporal occurrences in the context of thalamic functional impairments in patients with 22q11.2DS. Second, we touch upon the methodological innovations brought about by the proposed algorithm.

### Thalamic nuclei and functional correlates

The thalamus nuclei can be classified according to their function into three distinct groups: relay nuclei, association nuclei, and midline nuclei [55, 56]. The relay nuclei are responsible for relaying specific sensory information from the periphery to the cerebral cortex, while the association nuclei receive most of their information from the cerebral cortex and project back to the association areas in the cortex. Lastly, the midline nuclei have broad and diffuse projections through the cerebral cortex, and are involved in general functions such as consciousness and attention [57]. With our μCAP analysis, we were able to capture several of these functional networks in a data-driven manner. μCAP2 and μCAP3 both included strong midline thalamus signals. In agreement with past findings from network functional analysis [58], the DMN co-activated with the midline thalamus in both μCAPs 2 and 3. The midline thalamus is a group of nuclei that have been linked to a variety of cognitive functions, including emotion, memory, decision making, and attention [59]. It could be further divided into dorsal (paratenial and paraventricular nuclei) and ventral midline (reuniens and rhomboid nuclei) subcomponents [60].

Specifically, the ventral midline thalamus is thought to play a role in the integration of information from different brain regions and in the coordination of different cognitive processes [57, 61]. It has been suggested that the ventral midline thalamus acts as a bridge between different cortical regions, enabling communication between them, and that it is involved in the modulation of emotion and memory. Here, μCAP2 mostly included signal from the ventral midline thalamus and showed a pattern of co-activation of the DMN with the hippocampus and temporal regions. This finding is in line with studies linking the ventral midline thalamus with functional processes such as working memory, spatial navigation and contextual memory [62], as these processes are mediated by connections of the prefrontal cortex to the hippocampus [63]. In contrast, μCAP3 largely featured the dorsal midline thalamus, exhibited hippocampal deactivation and caudate, putamen and nucleus accumbens activations. These observations are reminiscent of studies linking the dorsal midline thalamus with feeding and drug seeking behaviors [55, 64].

μCAP4 revealed a pattern of co-activation between the mediodorsal thalamus and regions commonly associated with the salience and central executive networks, while negative signal was observed within the default mode network (DMN). The mediodorsal thalamus is considered an assortment of the associative nuclei which are believed to play a significant role in the function of the prefrontal cortex (PFC) [65]. Animal studies involving optogenetics and neurophysiology have demonstrated that the mediodorsal thalamus is responsible for thalamic mechanisms that contribute to cognitive functions carried out by the cortex. It is suggested that the mediodorsal thalamus optimizes connections between cortical neurons through a balance of excitation and feed-forward inhibition, which is necessary for cognitive switching and flexibility [23, 66, 67]. The co-activation of the executive and salience networks with the mediodorsal thalamus in μCAP4, concurrently with suppression of the DMN, provides further support for the role of the mediodorsal thalamus in cognitive switching and aligns with the triple network theory, which suggests that the suppression of the DMN is necessary for activation of the executive network [68].

### Patterns with clinical association in patients with 22q11.2DS

μCAP1 included contributions from several known functional circuitries. The lateral geniculate and medial geniculate nuclei are known to receive inputs from the superior and inferior colliculus, respectively, and as sensory relays, send outputs to the visual and auditory cortices [69, 70]. All aforementioned regions appeared active within μCAP1. Moreover, the lateral posterior nucleus of the thalamus (LP) receives inputs from the auditory, visual, and somatosensory cortices and plays a crucial role in the multi-sensory processing of information [71]. Additionally, in μCAP1, an isolated focus of activity was detected within the anteroventral thalamus along with the co-activation of limbic regions. Overall, this μCAP thus consists of multiple relay and associative nuclei within the thalamus with a complex pattern of multisensory cortical connections as well as limbic connections. We speculate that the co-occurrence of these circuits may aid integration within limbic and sensory processes. However, μCAP1 occurred significantly more in patients with 22q11.2DS. Thus, it is possible that hyper-occurrence of μCAP1 may contribute to disruption of the physiological gatekeeping role of the thalamus, possibly through hyper-recruitment of sensory circuits without higher-order control by the prefrontal regions. This is in agreement with the abnormal saliency model [29] of psychosis. Additionally the hyperconnectivity of thalamic-sensory regions is consistent with a previous study on auditory hallucinations in a partially overlapping sample, where hyperconnectivity between the medial geniculate nucleus/the anteroventral thalamus and the auditory cortex was detected [72]. Moreover, the recruitment of the hippocampus and midbrain dopaminergic regions observed in μCAP1 lends further evidence to the involvement of the thalamus in a priorly discussed hyper-dopaminergic state driven by hippocampal hyperactivity in patients with 22q11.2DS [73].

Additionally, regarding the involvement of the anteroventral thalamus in μCAP1, the hippocampus-anteroventral thalamus-prefrontal pathway is a known circuit for memory and temporal processing [74], an executive process that is impaired in patients with 22q11.2DS [75]. In fact, impairment of the hippocampus-thalamus-prefrontal circuits has been discussed as a possible biomarker of patients with 22q11.2DS [72, 76]. Consistently with this hypothesis, we found that prefrontal areas associated with this memory circuit, namely the dlPFC and dACC, were deactive in μCAP1. Prefrontal hypoactivity in patients with 22q11.2DS has been previously discussed and is reminiscent of recurrent findings in populations with psychosis spectrum. In particular, impaired thalamocortical connections are known to underlay prefrontal hypoactivity in patients with schizophrenia [77]. Additionally, prefrontal cortex dysconnectivity from the hippocampus and hippocampal-striatal hyperconnectivity are critical circuit impairments in patients with 22q11.2DS and contribute to psychosis susceptibility [73, 78]. The high occurrence of μCAP1 in these patients may be suggestive of a disconnection between the prefrontal cortex and the anteroventral thalamus within the PFC-anteroventral thalamus-hippocampus pathway.

The seed pattern within μCAP1 showed the highest similarity with that within μCAP6 (**Supplementary Materials, Figure S6**). Interestingly, like μCAP1, μCAP6 revealed a pattern of auditory, visual, and limbic regions. However, unlike μCAP1, the dlPFC and dACC were active within μCAP6, and μCAP6 occurred significantly more in HCs. Therefore, μCAP6 could be representative of a co-activation pattern that incorporates the recruitment of visuo-auditory circuits with the geniculate nuclei as well as the hippocampus-thalamus connection under the higher-order control of the dlPFC and dACC. Interestingly, the DMN was also deactive in μCAP6. DMN disengagement is observed during external and executive processing, and there is extensive evidence for the impairment of DMN disengagement in patients with 22q11.2DS [79, 80]. Taken together, our findings for μCAP1 and μCAP6 provide a dynamic framework for sensory pathway hyperactivity in 22q11.2DS. Accordingly, the co-occurrence of visuo-auditory circuits and executive regions, as well as the proper disengagement of the DMN, occur more in the healthy, whereas the shift towards the occurrence of visuo-auditory circuits without engaging executive cortices is a characteristic of patients with 22q11.2DS. This framework is well in line with evidence that shows preferential bias towards internally generated information in hallucinations (for a meta-analysis, see [81]), which may contribute to psychosis susceptibility in 22q11.2DS.

Finally, μCAP5 comprised activation of the mediodorsal thalamus along with both geniculate nuclei, which were co-activated with various cortical regions. The mediodorsal thalamus is composed of several distinct subdivisions, each with unique efferent and afferent connections to the prefrontal, cingulate, insular and supplementary motor cortices. Here, we found that this pattern of thalamo-cortical connectivity occurs significantly less in patients with 22q11.2DS. Moreover, within patients, μCAP5 occurred significantly less in patients with higher scores for positive psychotic symptoms. Thalamocortical dysconnectivity has been suggested to play a key role in the development of psychosis. Evidence suggests that thalamocortical connectivity is reduced in individuals suffering from psychosis and has been identified as a potential neural mechanism underlying the condition [82-84]. Neuroimaging studies have found decreased connectivity between the thalamus and the cortex in both recent-onset psychosis and schizophrenia. This dysfunction has been associated with altered perception, abnormal thinking, and bizarre behavior [85, 86].

Specifically, the mediodorsal thalamus-PFC connection is known to be essential for persistent PFC activity [87]. The mediodorsal thalamus is connected to the inhibitory interneurons within the PFC and is involved in allocating frontal cortex activity to the task at hand. Impairment of the mediodorsal thalamus-PFC circuit has been proposed in the pathogenesis of psychosis in that the disruption of the mediodorsal thalamus contributes to cognitive and behavioral deficits associated with psychosis [88, 89]. Furthermore, damage to the mediodorsal thalamus results in changes in the frontal cortex that account for some of the behavioral and cognitive deficits associated with psychosis [90]. Our results regarding the reduction of μCAP5 expression and its association with the severity of psychotic symptoms are well aligned with the notion of cortical dysconnectivity in psychosis. Moreover, the pattern of mediodorsal thalamus and geniculate activation within the thalamus points towards the two major roles that the thalamus may play in the pathophysiology of psychotic symptoms: sensory misattribution and aberrant modulation of prefrontal cortex. Crucially, this clinical association was only detectable when using the μCAP analysis that allowed for finer spatial information within the seed as well as a more robust temporal sampling along the time course of the RS-fMRI scans.

### Co-Activation Patterns and Methodological Contribution

One major contribution of the method that we devised was to allow for the detection of μCAPs such as μCAP2 and 6, where significant portions of the thalamus had negative values. In theory, the appearance of such patterns is solely possible within the framework of μCAP analysis, as the traditional approach averages the BOLD signal across the entire seed and therefore cancels out patterns where both positive and negative values are observable. Physiologically, co-occurrence of negative and positive values within the thalamus is particularly pertinent, as activation of one thalamic nucleus may lead to the inhibition of another mediated by the inhibitory sheet of nuclei (TRN) that encapsulates most of the thalamus [91, 92]. This allows the thalamus to process signals in a switch-board manner, either from sensory input or top-down control. Therefore, the activation pattern within μCAP2 and μCAP6 may be the result of an orchestrated excitation-inhibition arrangement within the nuclei that is crucial for thalamic functioning. Our simple toy model partly captures this feature of μCAPs.

Moreover, the μCAPs methodology allows to reveal contributions of functional units within the thalamus that are not mutually exclusive and may partly overlap. We believe that this feature is important for detecting dynamic functional circuits at the temporal resolution that fMRI provides. In fMRI, the sampling rate and the hemodynamic response are much slower than the rate of neuronal activity. Therefore, mutually exclusive functional parcels may not be as physiologically pertinent because it is imaginable that multiple sequential functional circuits appear together in one μCAP with functional relevance. For example, within μCAP5, the co-activation of the mediodorsal nucleus with geniculate nuclei represents an example of sensory and associative nuclei forming a functional unit together to contribute sensory input and top-down processing of the information.

As expected, intra-thalamic patterns detected by the μCAPs were spatially more diverse because this method allows for smaller parcels within the seed to be detected as “active”. Indeed, a more fine-grained definition of activation within the thalamus resulted in μCAPs sampling more functional frames across time to produce the co-activation patterns. Therefore, the whole-brain patterns produced by the μCAP algorithm were more spatially distinct from each other when compared with the traditional CAP methodology. Furthermore, retaining more temporal information with μCAP enhances the advantages of dynamic fMRI analysis in providing behaviorally relevant information.

### Limitations

The absence of cerebellum in the brain areas included into our analysis is a limitation of our data. Had cerebellar functional information been available in these scans, we believe that more networks involving the motor thalamus would have been resolved. Furthermore, while negative signal within the thalamus was discovered, our method so far only allowed for positive subparts to drive the activation patterns. Allowing negative subparts of the seed to contribute to the selection of frames under the notion of “deactive nuclei” may be explored in future investigations. In addition, we did not include individuals younger than 18 in the analysis. Future studies with a larger sample size are needed to investigate the effects of age and sex on functional patterns within the thalamus.

Finally, similar to previous studies in this cohort, the difference of head movement across patients with 22q11.2DS and HC remains a limitation in discussing the results regarding group comparison. To address this issue, during scrubbing, we opted for a low threshold (0.3 mm) to ensure overall low head movement. As a supplementary analysis, we regressed out movement and IQ as covariates of no interest in our group comparison to ensure that the results remain significant. The results of the analysis regarding head movement are depicted in the **Supplementary Materials**.

## Conclusion and perspectives

In sum, the proposed μCAP method is able to yield finer foci of activity within the initial seed region, which helps to retain valuable, and clinically relevant, spatiotemporal information. In particular, we showed that thalamic spatiotemporal dynamics differs across diagnostic groups, with relevance regarding psychopathology in the context of psychosis. In adults with 22q11.2DS, we discovered abnormally low occurrences of a μCAP including the mediodorsal thalamus and cortical connections, which were related to the severity of positive psychotic symptoms. Additionally, we revisited the evidence of geniculate hyperconnectivity with the audio-visual cortex in 22q11.2DS in the context of dynamic functional connectivity. In that regard, we showed that co-activation of geniculate nuclei with the audio-visual cortex is a pattern present in patients at rest with recruitment of the hippocampus and in the absence of higher-order cortical activity. On the other hand, in healthy controls, this circuit is more strongly recruited when the DMN is suppressed, and when higher-order prefrontal regions are active. Thus, our findings support and build upon existing research on the dysfunction of thalamic gatekeeping of sensory information in the pathophysiology of psychosis.

## Acknowledgments

This has been supported by the Swiss National Science Foundation (Grant Nos. 324730_121996 and 324730_144260 [to SE]) and a National Centre of Competence in Research Synapsy grant (Grant No. 51NF40-158776 [to SE]). We extend our deepest appreciation to all the families who actively participated in our research study. We would like to extend a special acknowledgment to Tereza Kotalova for her exceptional coordination of the project and to the MRI operators at Campus Biotech, namely Roberto Martuzzi, Loan Mattera, and Nathalie Philippe, for their valuable contributions. Lastly, we express our gratitude to Karin Bortolin for her generous assistance in conducting the statistical analysis.

## References

1. Ogawa, S., et al., Brain magnetic resonance imaging with contrast dependent on blood oxygenation. Proceedings of the National Academy of Sciences, 1990. 87(24): p. 9868–9872.

2. McGuire, P.K. and K. Matsumoto, Functional neuroimaging in mental disorders. World Psychiatry, 2004. 3(1): p. 6–11.

3. Zhan, X. and R. Yu, A Window into the Brain: Advances in Psychiatric fMRI. Biomed Res Int, 2015. 2015: p. 542467.

4. Allen, E.A., et al., Tracking whole-brain connectivity dynamics in the resting state. Cereb Cortex, 2014. 24(3): p. 663–76.

5. Bolton, T.A.W., et al., Interactions Between Large-Scale Functional Brain Networks are Captured by Sparse Coupled HMMs. IEEE Transactions on Medical Imaging, 2018. 37(1): p. 230–240.

6. Karahanoğlu, F.I. and D. Van De Ville, Transient brain activity disentangles fMRI resting-state dynamics in terms of spatially and temporally overlapping networks. Nature Communications, 2015. 6(1): p. 7751.

7. Vidaurre, D., S.M. Smith, and M.W. Woolrich, Brain network dynamics are hierarchically organized in time. Proceedings of the National Academy of Sciences, 2017. 114(48): p. 12827–12832.

8. Lurie, D.J., et al., Questions and controversies in the study of time-varying functional connectivity in resting fMRI. Netw Neurosci, 2020. 4(1): p. 30–69.

9. Preti, M.G., T.A. Bolton, and D. Van De Ville, The dynamic functional connectome: State-of-the-art and perspectives. Neuroimage, 2017. 160: p. 41–54.

10. Bolton, T.A.W., et al., Tapping into Multi-Faceted Human Behavior and Psychopathology Using fMRI Brain Dynamics. Trends in Neurosciences, 2020. 43(9): p. 667–680.

11. Canario, E., D. Chen, and B. Biswal, A review of resting-state fMRI and its use to examine psychiatric disorders. Psychoradiology, 2021. 1(1): p. 42–53.

12. Cole, D., S. Smith, and C. Beckmann, Advances and pitfalls in the analysis and interpretation of resting-state FMRI data. Frontiers in Systems Neuroscience, 2010. 4.

13. Chen, J.E., et al., Introducing co-activation pattern metrics to quantify spontaneous brain network dynamics. Neuroimage, 2015. 111: p. 476–88.

14. Liu, X. and J.H. Duyn, Time-varying functional network information extracted from brief instances of spontaneous brain activity. Proceedings of the National Academy of Sciences, 2013. 110(11): p. 4392–4397.

15. Liu, X., et al., Co-activation patterns in resting-state fMRI signals. Neuroimage, 2018. 180(Pt B): p. 485–494.

16. Bolton, T.A.W., et al., Triple Network Model Dynamically Revisited: Lower Salience Network State Switching in Pre-psychosis. Frontiers in Physiology, 2020. 11.

17. Ke, M., L. Hou, and G. Liu, The co-activation patterns of multiple brain regions in Juvenile Myoclonic Epilepsy. Cognitive Neurodynamics, 2022.

18. Rey, G., et al., Dynamics of amygdala connectivity in bipolar disorders: a longitudinal study across mood states. Neuropsychopharmacology, 2021. 46(9): p. 1693–1701.

19. Wang, L., et al., Deficient dynamics of prefrontal-striatal and striatal-default mode network (DMN) neural circuits in internet gaming disorder. J Affect Disord, 2023. 323: p. 336–344.

20. Dhanis, H., et al., Robotically-induced hallucination triggers subtle changes in brain network transitions. Neuroimage, 2022. 248: p. 118862.

21. Gaviria, J., et al., Dynamic functional brain networks underlying the temporal inertia of negative emotions. NeuroImage, 2021. 240: p. 118377.

22. Matsui, T., et al., On co-activation pattern analysis and non-stationarity of resting brain activity. NeuroImage, 2022. 249: p. 118904.

23. Zhou, K., et al., The Contribution of Thalamic Nuclei in Salience Processing. Frontiers in Behavioral Neuroscience, 2021. 15.

24. Hwang, K., et al., The Human Thalamus Is an Integrative Hub for Functional Brain Networks. J Neurosci, 2017. 37(23): p. 5594–5607.

25. Redinbaugh, M.J., et al., Thalamus Modulates Consciousness via Layer-Specific Control of Cortex. Neuron, 2020. 106(1): p. 66-75.e12.

26. Wang, S., et al., More Than Just Static: Dynamic Functional Connectivity Changes of the Thalamic Nuclei to Cortex in Parkinson’s Disease With Freezing of Gait. Frontiers in Neurology, 2021. 12.

27. Steullet, P., Thalamus-related anomalies as candidate mechanism-based biomarkers for psychosis. Schizophrenia Research, 2020. 226: p. 147–157.

28. Howes, O.D., et al., Aberrant Salience, Information Processing, and Dopaminergic Signaling in People at Clinical High Risk for Psychosis. Biological Psychiatry, 2020. 88(4): p. 304–314.

29. Roiser, J.P., et al., Neural and Behavioral Correlates of Aberrant Salience in Individuals at Risk for Psychosis. Schizophrenia Bulletin, 2012. 39(6): p. 1328–1336.

30. Knolle, F., et al., Brain responses to different types of salience in antipsychotic na ï ve first episode psychosis: An fMRI study. Translational Psychiatry, 2018. 8(1): p. 196.

31. Damaraju, E., et al., Dynamic functional connectivity analysis reveals transient states of dysconnectivity in schizophrenia. NeuroImage: Clinical, 2014. 5: p. 298–308.

32. Çetin, M.S., et al., Thalamus and posterior temporal lobe show greater inter-network connectivity at rest and across sensory paradigms in schizophrenia. NeuroImage, 2014. 97: p. 117–126.

33. Herrero, M.T., C. Barcia, and J.M. Navarro, Functional anatomy of thalamus and basal ganglia. Childs Nerv Syst, 2002. 18(8): p. 386–404.

34. Tregidgo, H.F.J., et al., Accurate Bayesian segmentation of thalamic nuclei using diffusion MRI and an improved histological atlas. NeuroImage, 2023. 274: p. 120129.

35. Bezdudnaya, T. and A. Keller, Laterodorsal nucleus of the thalamus: A processor of somatosensory inputs. J Comp Neurol, 2008. 507(6): p. 1979–89.

36. Krause, T., et al., Thalamic sensory strokes with and without pain: differences in lesion patterns in the ventral posterior thalamus. Journal of Neurology, Neurosurgery &amp; Psychiatry, 2012. 83(8): p. 776–784.

37. Saalmann, Y.B., Intralaminar and medial thalamic influence on cortical synchrony, information transmission and cognition. Frontiers in Systems Neuroscience, 2014. 8.

38. Jiang, Y., M.H. Patton, and S.S. Zakharenko, A Case for Thalamic Mechanisms of Schizophrenia: Perspective From Modeling 22q11.2 Deletion Syndrome. Frontiers in Neural Circuits, 2021. 15.

39. Hwang, K., et al., Neuropsychological evidence of multi-domain network hubs in the human thalamus. eLife, 2021. 10: p. e69480.

40. Óskarsdóttir, S., et al., Updated clinical practice recommendations for managing children with 22q11.2 deletion syndrome. Genetics in Medicine, 2023. 25(3): p. 100338.

41. Schneider, M., et al., Clinical and cognitive risk factors for psychotic symptoms in 22q11.2 deletion syndrome: a transversal and longitudinal approach. European Child & Adolescent Psychiatry, 2014. 23(6): p. 425–436.

42. Bassett, A.S. and E.W. Chow, 22q11 deletion syndrome: a genetic subtype of schizophrenia. Biological psychiatry, 1999. 46(7): p. 882–891.

43. First, M.B., et al., Structured clinical interview for DSM-IV-TR Axis I disorders: patient edition. 2005: Biometrics Research Department, Columbia University New York, NY.

44. Kay, S.R., A. Fiszbein, and L.A. Opler, The positive and negative syndrome scale (PANSS) for schizophrenia. Schizophrenia bulletin, 1987. 13(2): p. 261–276.

45. Zöller, D., et al., Disentangling resting-state BOLD variability and PCC functional connectivity in 22q11.2 deletion syndrome. NeuroImage, 2017. 149: p. 85–97.

46. Yan, C. and Y. Zang, DPARSF: a MATLAB toolbox for” pipeline” data analysis of resting-state fMRI. Frontiers in systems neuroscience, 2010: p. 13.

47. Alemán-Gómez, Y. IBASPM: toolbox for automatic parcellation of brain structures. in 12th Annual Meeting of the Organization for Human Brain Mapping. June 11-15, 2006. Florence, Italy. 2006.

48. Ashburner, J. and K.J. Friston, Unified segmentation. Neuroimage, 2005. 26(3): p. 839–851.

49. Ashburner, J., A fast diffeomorphic image registration algorithm. Neuroimage, 2007. 38(1): p. 95–113.

50. Power, J.D., et al., Spurious but systematic correlations in functional connectivity MRI networks arise from subject motion. Neuroimage, 2012. 59(3): p. 2142–54.

51. Bolton, T.A.W., et al., TbCAPs: A toolbox for co-activation pattern analysis. NeuroImage, 2020. 211: p. 116621.

52. Kuhn, H.W., Variants of the hungarian method for assignment problems. Naval Research Logistics Quarterly, 1956. 3(4): p. 253–258.

53. Tzourio-Mazoyer, N., et al., Automated anatomical labeling of activations in SPM using a macroscopic anatomical parcellation of the MNI MRI single-subject brain. Neuroimage, 2002. 15(1): p. 273–89.

54. Benjamini, Y. and Y. Hochberg, Controlling the False Discovery Rate: A Practical and Powerful Approach to Multiple Testing. Journal of the Royal Statistical Society: Series B (Methodological), 1995. 57(1): p. 289–300.

55. Groenewegen, H.J. and M.P. Witter, CHAPTER 17 - Thalamus, in The Rat Nervous System (Third Edition), G. Paxinos, Editor. 2004, Academic Press: Burlington. p. 407-453.

56. Iglesias, J.E., et al., A probabilistic atlas of the human thalamic nuclei combining ex vivo MRI and histology. NeuroImage, 2018. 183: p. 314–326.

57. Reeders, P.C., et al., Identifying the midline thalamus in humans <em>in vivo</em>. bioRxiv, 2022: p. 2022.02.20.481099.

58. Kumar, V.J., et al., Relay and higher-order thalamic nuclei show an intertwined functional association with cortical-networks. Communications Biology, 2022. 5(1): p. 1187.

59. Vertes, R.P., S.B. Linley, and A.K.P. Rojas, Structural and functional organization of the midline and intralaminar nuclei of the thalamus. Front Behav Neurosci, 2022. 16: p. 964644.

60. Ding, S.L., et al., Comprehensive cellular-resolution atlas of the adult human brain. J Comp Neurol, 2016. 524(16): p. 3127–481.

61. Vertes, R.P., Differential projections of the infralimbic and prelimbic cortex in the rat. Synapse, 2004. 51(1): p. 32–58.

62. Vertes, R.P., S.B. Linley, and W.B. Hoover, Limbic circuitry of the midline thalamus. Neurosci Biobehav Rev, 2015. 54: p. 89–107.

63. Jung, D., Y. Huh, and J. Cho, The Ventral Midline Thalamus Mediates Hippocampal Spatial Information Processes upon Spatial Cue Changes. The Journal of Neuroscience, 2019. 39(12): p. 2276–2290.

64. Li, S. and G.J. Kirouac, Sources of inputs to the anterior and posterior aspects of the paraventricular nucleus of the thalamus. Brain Structure and Function, 2012. 217: p. 257–273.

65. Nakajima, M. and M.M. Halassa, Thalamic control of functional cortical connectivity. Curr Opin Neurobiol, 2017. 44: p. 127–131.

66. Rikhye, R.V., A. Gilra, and M.M. Halassa, Thalamic regulation of switching between cortical representations enables cognitive flexibility. Nature Neuroscience, 2018. 21(12): p. 1753–1763.

67. Larsen, R., et al., The Thalamus Regulates Retinoic Acid Signaling and Development of Parvalbumin Interneurons in Postnatal Mouse Prefrontal Cortex. eNeuro, 2019. 6(1).

68. Anticevic, A., et al., The role of default network deactivation in cognition and disease. Trends in Cognitive Sciences, 2012. 16(12): p. 584–592.

69. Winer, J.A., The functional architecture of the medial geniculate body and the primary auditory cortex. The mammalian auditory pathway: Neuroanatomy, 1992: p. 222–409.

70. Preston, A. and A.S. Evans, Lateral Geniculate Nucleus of Thalamus, in Encyclopedia of Clinical Neuropsychology, J.S. Kreutzer, J. DeLuca, and B. Caplan, Editors. 2011, Springer New York: New York, NY. p. 1435-1436.

71. Zhang, W. and R.M. Bruno, High-order thalamic inputs to primary somatosensory cortex are stronger and longer lasting than cortical inputs. eLife, 2019. 8: p. e44158.

72. Mancini, V., et al., Abnormal Development and Dysconnectivity of Distinct Thalamic Nuclei in Patients With 22q11.2 Deletion Syndrome Experiencing Auditory Hallucinations. Biological Psychiatry: Cognitive Neuroscience and Neuroimaging, 2020. 5(9): p. 875–890.

73. Delavari, F., et al., Dysmaturation Observed as Altered Hippocampal Functional Connectivity at Rest Is Associated With the Emergence of Positive Psychotic Symptoms in Patients With 22q11 Deletion Syndrome. Biol Psychiatry, 2021. 90(1): p. 58–68.

74. Nelson, A.J.D., The anterior thalamic nuclei and cognition: A role beyond space? Neuroscience & Biobehavioral Reviews, 2021. 126: p. 1–11.

75. Maeder, J., et al., From Learning to Memory: A Comparison Between Verbal and Non-verbal Skills in 22q11.2 Deletion Syndrome. Frontiers in Psychiatry, 2021. 12.

76. Schleifer, C., et al., Dissociable Disruptions in Thalamic and Hippocampal Resting-State Functional Connectivity in Youth with 22q11.2 Deletions. J Neurosci, 2019. 39(7): p. 1301–1319.

77. Popken, G.J., et al., Subnucleus-specific loss of neurons in medial thalamus of schizophrenics. Proc Natl Acad Sci U S A, 2000. 97(16): p. 9276–80.

78. Mukherjee, A., et al., Long-Lasting Rescue of Network and Cognitive Dysfunction in a Genetic Schizophrenia Model. Cell, 2019. 178(6): p. 1387-1402.e14.

79. Padula, M.C., et al., Structural and functional connectivity in the default mode network in 22q11.2 deletion syndrome. J Neurodev Disord, 2015. 7(1): p. 23.

80. Dubourg, L., et al., Divergent default mode network connectivity during social perception in 22q11.2 deletion syndrome. Psychiatry Research: Neuroimaging, 2019. 291: p. 9–17.

81. Kompus, K., R. Westerhausen, and K. Hugdahl, The “paradoxical” engagement of the primary auditory cortex in patients with auditory verbal hallucinations: a meta-analysis of functional neuroimaging studies. Neuropsychologia, 2011. 49(12): p. 3361–9.

82. Harrison, P.J., The neuropathology of schizophrenia: a critical review of the data and their interpretation. Brain, 1999. 122(4): p. 593–624.

83. Hazlett, E.A., et al., Three-dimensional analysis with MRI and PET of the size, shape, and function of the thalamus in the schizophrenia spectrum. American Journal of Psychiatry, 1999. 156(8): p. 1190–1199.

84. Avram, M., et al., Bridging the Gap? Altered Thalamocortical Connectivity in Psychotic and Psychedelic States. Frontiers in Psychiatry, 2021. 12.

85. Ramsay, I.S., et al., Thalamocortical connectivity and its relationship with symptoms and cognition across the psychosis continuum. Psychol Med, 2022: p. 1–10.

86. Yao, B., et al., Altered thalamocortical structural connectivity in persons with schizophrenia and healthy siblings. Neuroimage Clin, 2020. 28: p. 102370.

87. Pergola, G., et al., The Regulatory Role of the Human Mediodorsal Thalamus. Trends Cogn Sci, 2018. 22(11): p. 1011–1025.

88. Popken, G.J., et al., Subnucleus-specific loss of neurons in medial thalamus of schizophrenics. Proceedings of the National Academy of Sciences, 2000. 97(16): p. 9276–9280.

89. Skåtun, K.C., et al., Thalamo-cortical functional connectivity in schizophrenia and bipolar disorder. Brain Imaging Behav, 2018. 12(3): p. 640–652.

90. Parnaudeau, S., S.S. Bolkan, and C. Kellendonk, The Mediodorsal Thalamus: An Essential Partner of the Prefrontal Cortex for Cognition. Biol Psychiatry, 2018. 83(8): p. 648–656.

91. Crabtree, J.W. and J.T.R. Isaac, New Intrathalamic Pathways Allowing Modality-Related and Cross-Modality Switching in the Dorsal Thalamus. The Journal of Neuroscience, 2002. 22(19): p. 8754.

92. Crabtree, J.W., Functional Diversity of Thalamic Reticular Subnetworks. Frontiers in Systems Neuroscience, 2018. 12.

