## Supplementary Materials for "Thalamic contributions to psychosis susceptibility: Evidence from co-activation patterns accounting for intra-seed spatial variability (μCAPs)"

**Methods**

*Validation with simulated data*

To validate and illustrate the benefits of µCAPs over traditional CAPs, we generated simulated data with characteristic spatiotemporal properties (**Figure S1**). One out of six possible states of activation was assumed to be expressed at each time point. Frames included ten voxels, of which two (voxels 1 and 6) constituted the seed region. Seed activity was only homogeneous in states 5 and 6, so that the ability to properly retrieve CAPs in more complex cases could be ascertained. In states 1 and 3, only half of the seed region was active, while in states 2 and 4, half of the seed region was active while the other was deactive. Simulated data were created with added independent Gaussian noise of standard deviation 0.1 (low noise setting) or 0.75 (high noise setting). Conventional CAPs were extracted with *TbCAPs*, and compared to µCAPs in terms of spatial patterns (quantifying the cosine distance between ground truth and extracted states) and transition probabilities (1 minus Pearson correlation between the ground truth and extracted transition probability matrices). For this toy example, a frame selection threshold of 0.5 was always used across iterations and methods.

At a low noise regime, µCAPs perfectly matched the ground truth state patterns (**Figure S1c**, top panel), while standard CAPs did not (**Figure S1c**, middle panel). More specifically, in this latter case, only patterns of activation were retrieved within the seed region. Similarly, transition probabilities were closely approximated when deriving µCAPs, but not standard CAPs, for which convergence onto the baseline state (S6) was almost exclusively captured **(Figure S1c**, right column). When noise was increased to a level where state transitions could not be visually discerned anymore (**Figure S1b**, right panel), extracted µCAPs still reasonably matched ground truth state patterns **(Figure S1c**, bottom). The correct transition probability pattern was also still recovered for transitions between states 1 to 5, although erroneous returns to baseline also started being estimated (**Figure S1c**, bottom right panel). In short, µCAPs can thus reveal spatiotemporal state features to which standard CAPs are insensitive, even in more realistic high-noise settings.

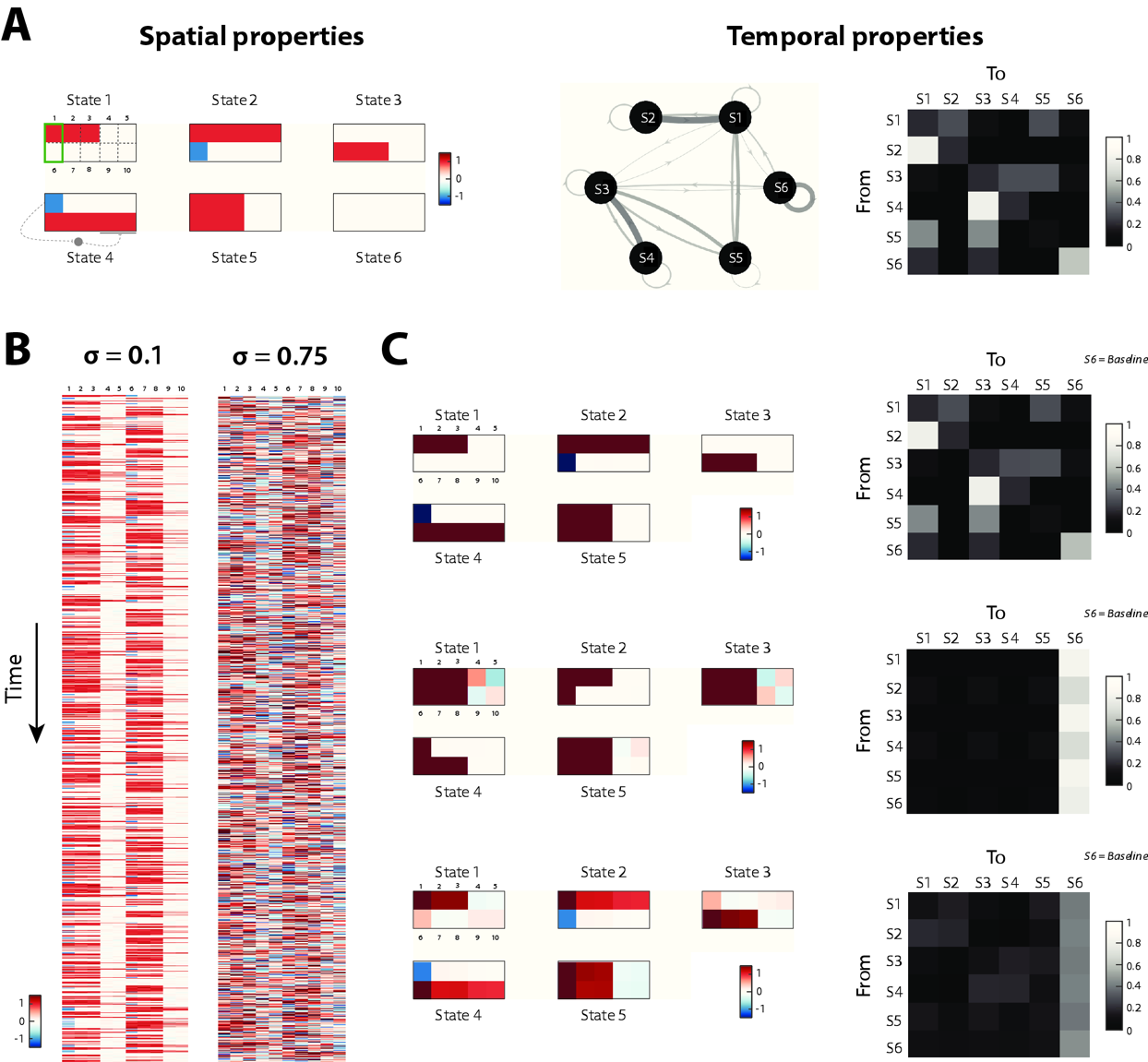

**Figure S1 - Results on simulated data.** (**a**) Spatial (left) and temporal (right) properties of the six considered states. States are made of ten voxels, with the two seed voxels highlighted by a green rectangle. (**b**) Example voxel-wise time courses of activity at a low noise (left) or high noise (right) regime. In the latter case, structured patterns cannot be discerned anymore. (**c**) Estimated spatial patterns (left) and transition probabilities (right) for µCAPs with low noise (top row), CAPs with low noise (middle row) or µCAPs with high noise (bottom row).

*Selection of optimal number of clusters*

The optimal number of clusters for K-means clustering was chosen using 2 different criteria based on a test – training process. For the first criterion, a 20-fold random split of subjects into half was done, and both test and training subjects went through frame selection. The training frames went through K-means clustering to produce the *K* different centroids (training centroids). Test frames were assigned to the training centroids from which they had the lowest distance. This distance was calculated as (1-correlation) to match the distance used within the K-means algorithm. The cluster with the highest sum of distances of test frames was chosen as the worst-fit cluster and it was monitored across a range of candidate *K* values (2 to 10).

As expected, the sum of distances decreased as *K* increased (**Figure S2a**). However, an optimum was indicated by a slower decrease of the sum of distances from *K*=6 onwards.

In the second criterion, another 20-fold random split of subjects into half was done and both test and training subjects went through frame selection. Here, both test and training frames went through K-means clustering to produce *K* different training and test centroids. Then, the *K* test centroids and *K* training centroids were matched using the Hungarian algorithm [50]. The average of the *K* distances across the matched centroids was calculated for a range of candidate *K* values (4 to 8). The investigated range was smaller owing to the heavy computational burden. The lowest distance of matched centroids would indicate maximal similarity of results across test and training subjects and therefore, stable K-means results.

In our experimental data, *K*=6 was once again found as an optimal solution (**Figure S2b**).

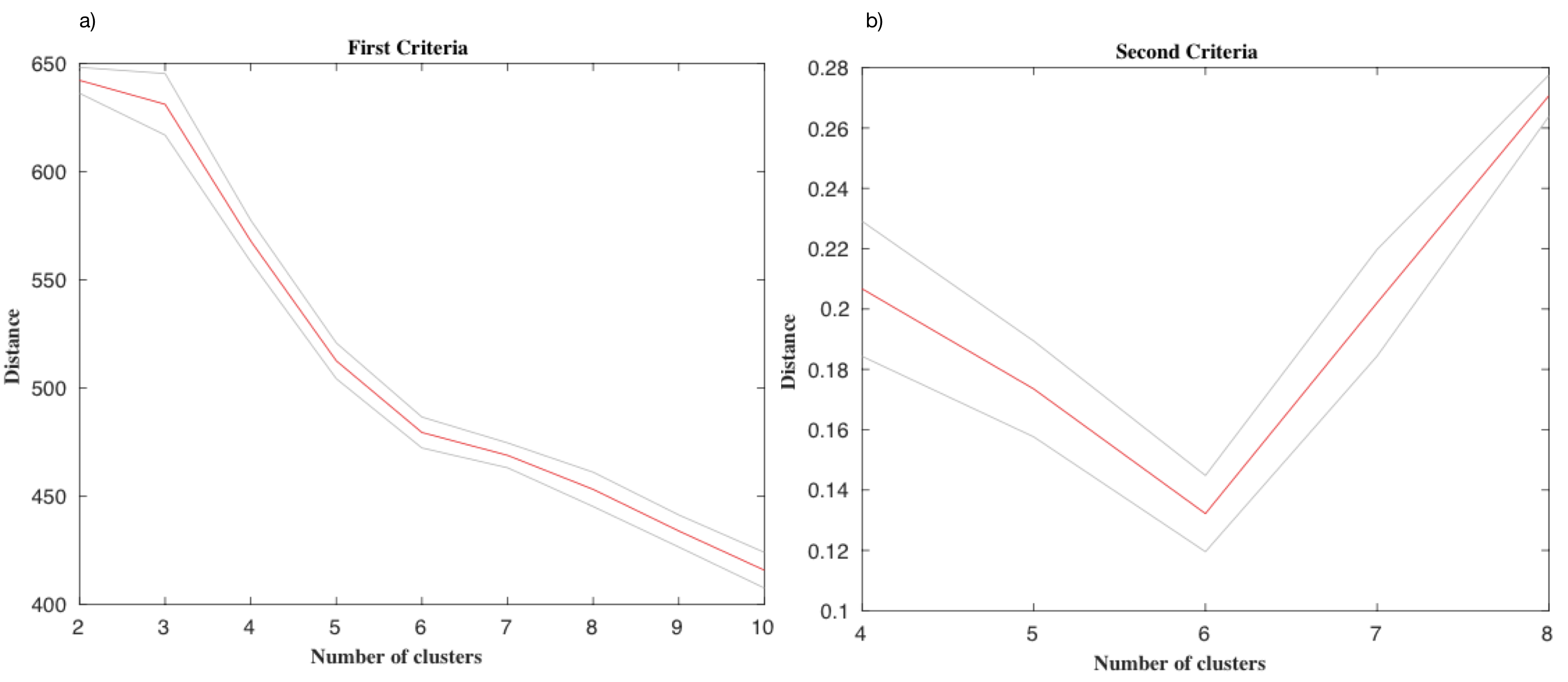

**Figure S2 - K=6 is an optimal number of clusters as seen from two distinct criteria.** (**a**) Sum of distances of test frames for the worst-fit cluster for each candidate K vale within the range of 2 to 10. K=6 was chosen as the optimal value. (**b**) Average distance of K centroids matched across test and training subjects for candidate K values within the range of 4 to 8. K=6 provided the minimum distance and therefore, the most stable K-means results. Results are displayed as average and 95% confidence interval over a 20-fold random split of subjects into half.

*Convergence assessment*

Convergence is monitored by keeping track of the distance between the seed pattern within a µCAP across iterations. First, µCAPs are matched across successive iterations using the Hungarian algorithm [50]. Then, the average pairwise cosine distance across µCAPs is computed. The threshold for convergence was set at 0.005. The initial seed pattern (here containing the entire thalamus) was left out of the convergence assessment.

**Figure S3** displays average distances over 15 consecutive iterations. Convergence can clearly be observed from the fifth iteration onwards.

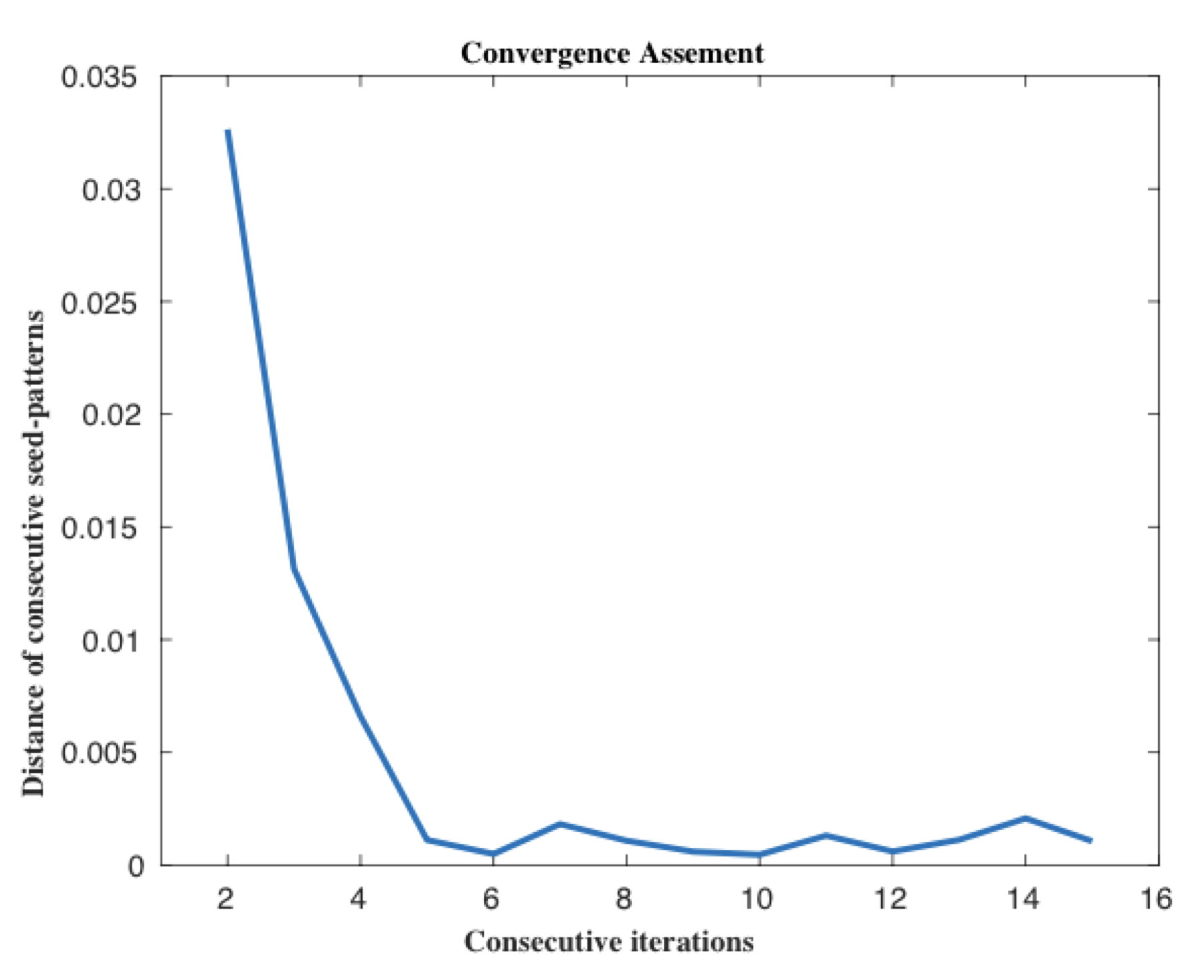

**Figure S3 - Distance between seed patterns within µCAPs across subsequent iterations.** The distance value dropped below the convergence threshold when comparing the seed patterns of iterations 4 and 5. The seed patterns of the first and second iterations are not compared, as the first seed pattern was the whole thalamus.

**Results**

*Six co-activation patterns driven by CAPs*

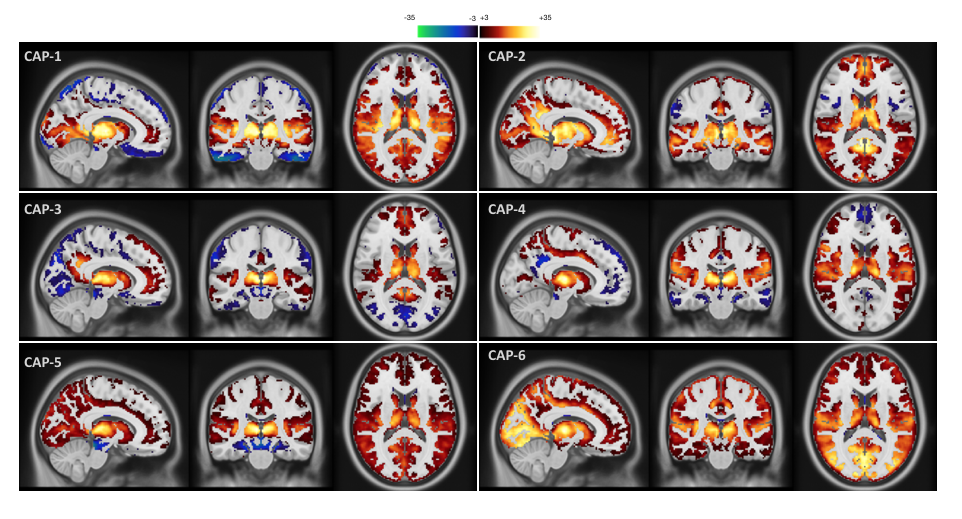

**Figure S4 – Conventional co-activation patterns extracted from the analyzed dataset.** Spatial T-score maps are calculated, for each voxel, as the average value for frames assigned to each co-activation pattern (CAP) divided by the standard deviation of the voxels across those frames multiplied by the square root of the number of frames. The results are thresholded at 3, corresponding to p < 0.01.

*Movement analysis*

**Table S1 – Impact of head movement on the results.** Frame-wise displacement across both clinical groups is shown before and after scrubbing (top). Then, analytical results (association of (µ)CAP occurrences with diagnosis and positive psychotic symptoms) are shown when frame-wise displacement after scrubbing and IQ were regressed out of the occurrences as covariates of no interest. Bold font denotes significant outcomes (^*^: p<0.05, ^**^: p<0.005, ^***^: p<0.001).

| **Variable Name** | | **22q11.2DS** | | | **HC** | | ***P*-value** | |
| --- | --- | --- | --- | --- | --- | --- | --- | --- |
| Frame-wise displacement before scrubbing [mean (SD)) | | 0.23 (0.07) | | | 0.17 (0.06) | | **<0.001 ***** | |
| Frame-wise displacement after scrubbing [mean (SD)) | | 0.14 (0.03) | | | 0.11 (0.03) | | **<0.001***** | |
| µCAPs | | | | | | | | |
|  | Association with Diagnosis | | | | | Association with Positive Psychotic Symptoms | | |
| µCAP number | Average occurrence regressed (SD) | | | *P*-value  (Wilcoxon rank sum test) | | Spearman’s correlation coefficient | | *P*-value |
|  | 22q11DS | | HC |  | | | | |
| 1 | 1.8148 | | -1.6204 | **0.0019 *** | | 0.1357 | | 0.3474 |
| 2 | -0.2480 | | 0.2215 | 0.3982 | | -0.0381 | | 0.7929 |
| 3 | -0.2396 | | 0.2140 | 0.6464 | | -0.0966 | | 0.5044 |
| 4 | -0.3654 | | 0.3262 | 0.3473 | | -0.1713 | | 0.2343 |
| 5 | -1.8380 | | 1.6411 | **0.0274 *** | | -0.3494 | | **0.0129 *** |
| 6 | -1.0361 | | 0.9251 | **0.0121 *** | | 0.2738 | | 0.0544 |
| Standard CAPs | | | | | | | | |
|  | Association with Diagnosis | | | | | Association with Positive Psychotic Symptoms | | |
| CAP number | Average occurrence regressed (SD) | | | *P*-value  (Wilcoxon rank sum test) | | Spearman’s correlation coefficient | | *P*-value |
|  | 22q11DS | | HC |  | | | | |
| 1 | 1.0943 | | -0.9771 | **0.0009 **** | | 0.0862 | | 0.5517 |
| 2 | -0.4344 | | 0.3878 | 0.0623 | | 0.1359 | | 0.3466 |
| 3 | -0.3748 | | 0.3346 | 0.3571 | | 0.0047 | | 0.9743 |
| 4 | -0.3135 | | 0.2799 | 0.2354 | | -0.0456 | | 0.7531 |
| 5 | -0.2699 | | 0.2410 | 0.1187 | | -0.1994 | | 0.1650 |
| 6 | -1.0883 | | 0.9717 | **0.0025 **** | | -0.1674 | | 0.2454 |

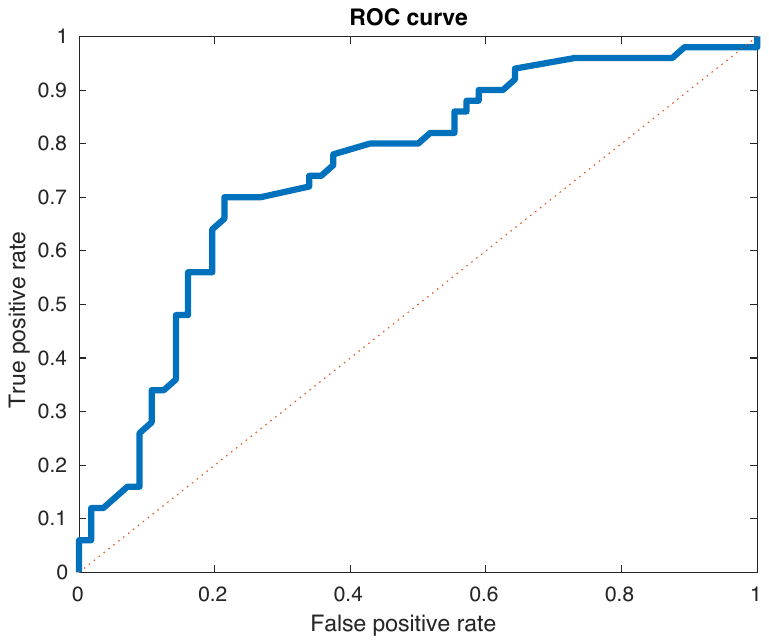

**Figure S5 - Receiver operating characteristic (ROC) curve for the detection of patients with 22q11.2DS.** An area under the curve (AUC) of 0.75 was found. µCAP1 and µCAP6 had similar within-seed patterns (r=0.68) but were expressed in different directions across healthy controls and patients with 22q11.2DS. Thus, we performed a post-hoc analysis on the occurrence ratio (µCAP6 over µCAP1). A significant group difference was found (Wilcoxon rank test, p-value<0.001).

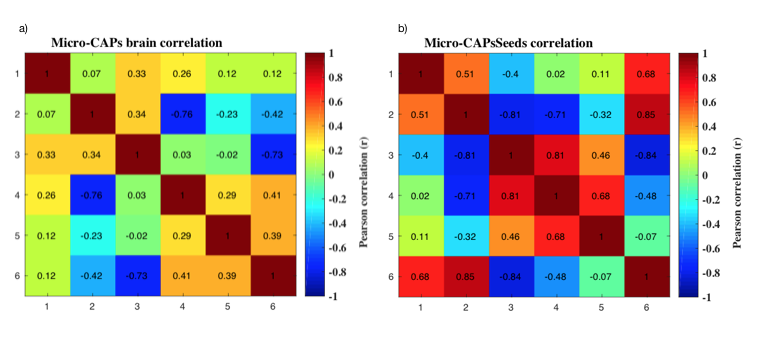

**Figure S6 – Pattern similarity within µCAPs.** Pairwise spatial correlations between co-activation patterns obtained using the µCAPs algorithm are displayed for whole-brain patterns (left) and within-seed patterns (right).
